# Timing of water limitations affects source to sink differences in δ^13^C composition in apple

**DOI:** 10.1101/2021.07.20.453133

**Authors:** Lee Kalcsits, Nadia Valverdi, Michelle Reid

**Affiliations:** Tree Fruit Research and Extension Center, Washington State University, Wenatchee, WA, United States; Department of Horticulture, Washington State University, Pullman, WA, United States

**Keywords:** fruit, leaves, roots, stems, carbon isotope, deficit irrigation

## Abstract

Deficit irrigation is used to reduce vegetative vigor, increase fruit quality, and conserve water resources. However, physiological responses to deficit irrigation can vary depending on soil and environmental conditions. Although physiological measurements are often made at single points in time, responses are often longer lasting and a measurement that integrates responses over time would have greater value in assessing the effectiveness of deficit irrigation practices. Carbon isotope composition has long been used as a proxy measurement for water-use efficiency, stomatal conductance, and carbon dioxide exchange with the atmosphere and is heavily influenced by water status. Potentially, fruit, leaves, or other tissues could be used as samples for carbon isotope measurements. However, it is not well known how irrigation practices can influence both source and sink tissue carbon isotope composition in perennial systems. Here, we used two experiments to determine how irrigation timing affects both source and sink δ^13^C at the end of the season. Irrigation limitations were initiated after bloom for either the whole season or for early, middle, or late season and compared to a well-watered control. For both experiments, leaves were poor indicators of irrigation deficit treatments that were applied during the season. There were no significant differences in leaf δ^13^C between deficit treatments and the control for both experiments. However, all sink tissues including roots and stems in experiment one for both years and for fruits in experiment two for both years were significantly more enriched compared to the well-watered control. Environmental conditions during the season also appeared to influence the magnitude of difference in δ^13^C between deficit irrigation treatments and the control. These results indicate that the use of sink tissues are more sensitive for measuring signals associated with in-season water deficits. Carbon isotope composition can be an effective proxy to measure efficacy of irrigation treatments at the physiological level.

## INTRODUCTION

Irrigation efficiency for fruit production is important to control vegetative vigor, optimize fruit quality (Talluto et al. 2008), and to conserve water resources (Einhorn and Caspari 2003; Fereres and Soriano 2006). There is interest in using deficit irrigation as a tool to control fruit size and to improve fruit quality in Honeycrisp apple. The adoption of deficit irrigation in most apple cultivars has been limited because of concerns of reductions in fruit size and overall plant yield. However, for Honeycrisp apple, fruit size is frequently large, and can contribute to high economic losses from postharvest disorders such as bitter pit (Cheng and Sazo 2018).

Carbon isotope discrimination is an effective proxy for intrinsic water-use efficiency and is primarily a function of carbon fluxes between the air and intercellular space where lower stomatal conductance will lead to greater increase δ^13^C of fixed carbon in the leaf (Farquhar et al. 1989). In theory, carbon isotope composition is useful to measure time-averaged plant water status. However, for perennial plants that regularly recycle and remobilize carbon to sink tissues at different times during the season (Furze et al. 2018), δ^13^C values can become difficult to interpret because of mixing of current and past season carbon pools. Similarly, this can also apply to short-term deficit treatments where carbon acquired prior to starting treatments can dilute the observed changes in carbon isotope composition of carbon acquired after deficit irrigation was initiated.

The timing of carbon fixation and transport to sink tissues is important to consider when measuring different plant tissues for carbon isotope composition. For example, for leaf formation, much of the structural carbon allocated during leaf development occurs early in the season (Marchi et al. 2005) and carbon acquired by the leaf afterwards is transported to sink tissues such as fruit, new growth, roots or stems. Here, we measured the δ^13^C of bulk tissue in apple during two deficit irrigation experiments. One experiment was focused on differences between both stem and root tissue compared to leaf tissue during season long deficit irrigation. The second experiment focused on how timing of deficit treatments during the season affected differences in carbon isotope composition between leaves and fruit. The hypothesis was that changes in the δ^13^C signal would be affected by the timing of carbon allocation to different plant tissues.

## MATERIALS AND METHODS

### Experiment 1: Leaf, root, and stem δ^13^C under season-long post-bloom deficit irrigation

The first experiment was conducted in a ‘Honeycrisp’ orchard that was grafted on M9-T337 and planted in 2016 at WSU-TFREC Sunrise Orchard on a shallow sandy loam soil and trained on a slender spindle system at a spacing of 0.9 m between trees and 3.6 m between rows. In 2017 and 2018, trees were drip irrigated daily using emitters spaced 30 cm apart that applied 3.78 L h^−1^ water for two hours (four periods of 30 minutes) daily and were fertilized using standard commercial practices. Each plot had five trees where the outer trees acted as border trees using the inner three trees for measurements.

The treatments consisted of a drought stress treatment maintaining soil water content near 50% FC and a well-watered control with irrigation maintaining soil water content near 100% FC. Field capacity was estimated to be approximately 33% vol/vol. In the field, one capacitance sensor was added per replication(N=3) and were installed at the beginning of the season (April) to record volumetric water content (m^3^/m^3^) every 30 min using Em50G 2.24 sensors (Decagon Devices, Pullman, WA, USA). Irrigation treatments were initiated 30 days after full bloom and maintained for 90 days. Irrigation decisions were soil-moisture based and once soil moisture had been depleted to 50% of field capacity, irrigation was applied to achieve soil moisture above 60% of field capacity. At the end of the experiment in 2018, whole trees were removed from the orchard and were separated into roots, stems and leaves. Roots were carefully washed using tap water to remove soil residues. Samples were then dried at 60 °C for seven days and stems were dried for 30 days.

### Experiment 2: Leaf and fruit δ^13^C under periodic deficit irrigation

The second experiment was also conducted at the WSU Sunrise Research Orchard located in Rock Island, WA using 240 three-year-old ‘Honeycrisp’ trees grafted on M9-T337 rootstock that were planted in 2015 and trained to a spindle training system. The planting density was 0.9 x 3.6 m. The soil is an alluvial shallow sandy loam soil. The first-year crop was in 2017. Using a completely randomized design, irrigation regimes were used that withheld irrigation either early, middle or late in the season and were compared to a fully watered control. Each replicate consisted of 20 trees with a total of 3 replicates per treatment. Water was applied to the well-watered control in amounts that exceeded water demand with a daily schedule of four 30-minute applications each day. The early season irrigation deficit was from 15 to 45 DAFB, mid-season irrigation deficit was from 45 to 75 DAFB and late season irrigation deficit was from 75 to105 DAFB. All treatments followed the well-watered irrigation schedule of the control before and after each deficit period. Soil volumetric water content and soil temperature were measured with an ECH2O 5TM soil moisture and temperature probes (Decagon Devices, Pullman, WA, USA) placed in each replicate at 20 cm depth in the herbicide strip directly between trees. Each probe was interfaced with an EM50G cellular data logger (Decagon Devices, Pullman, WA, USA) and data was logged every 30 minutes from full bloom to harvest. The deficit irrigation target was approximately 30-40% of field capacity and the well-watered control was approximately 85-100% of field capacity. During deficit periods, when the volumetric water content fell below 30% of field capacity, water was applied in small amounts to ensure that soil volumetric water content did not exceed 50% of field capacity. During these small irrigation sets, deficit trees only received four applications of 30 min each during a four-hour period. At the end of the season, fruit and leaf tissue samples from harvest were both used for carbon isotope analysis. Fruit was prepared for isotope analysis by slicing samples into thin segments and removing the seeds, peel and core and dried for seven days at 60 °C.

### Carbon Isotope Sampling

Leaf and fruit samples were ground into a fine powder using a VWR high throughput homogenizer (VWR, Radnor, PA, USA). For stems and roots, samples were initially ground to 20-micron size using a Wiley Mini mill (Thomas Scientific LLC., Swedesboro, NJ, USA) and then ground to sub-micron size using a VWR high throughput homogenizer (VWR, Radnor, PA, USA). Then samples of 3 μg for roots and leaf and 10 μg for stems and fruit were sent for δ^13^C composition using a using an Costech EXS4010 Elemental Analyzer (Costech Analytical, Valencia, California, USA) coupled with a ThermoFinniganDela Plus XP isotope ratio mass spectrometer (Thermo Electron, Bremen, Germany) at the Washington State University Stable Isotope Core Laboratory (SICL) in Pullman, WA. Stable isotope ratios were calculated using the following equation:

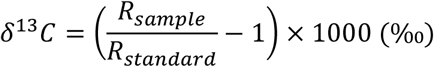

Where R_sample_ is the ^13^C/^12^C ratio of the sample and R_standard_ is the ^13^C/^12^C ratio of Vienna Pee Dee Belemnite calcium carbonate.

### Environmental Data

Environmental data including air temperature, relative humidity, wind speed, total precipitation, and solar radiation were obtained through the AgWeatherNet weather stations located at the WSU-Sunrise Orchard and WSU-TFREC. Data were analyzed by performing an analysis of variance (ANOVA), and a Tukey’s means separation with a confidence of 95% (SAS, ver. 9.4 PROC GLM).

## RESULTS AND DISCUSSION

### Experiment 1

Total tree dry weight was approximately 25% lower when deficit irrigation was applied compared to the well-watered control (P<0.05; Fig. 1). Leaf dry mass was less affected by deficit irrigation and was 20% lower than the control. However, root and stem biomass was approximately 25% lower compared to the control. Leaf δ^13^C for deficit irrigated trees was not significantly different than the control. Similar results were reported by Van Hooijdonk et al. (2004) in ‘Pacific Rose’ apple where deficit irrigation did not induce a change in δ^13^C of leaves. However, both root and stem δ^13^C was significantly greater than the well-watered control. When trees were well-watered, there were no significant differences among plant tissues in δ^13^C composition. However, when water was limited after bloom, root and stem δ^13^C composition was up to 3‰ different than the well-watered control. Greater differences were observed in 2017 than 2018 mostly because of greater δ^13^C in the control plant tissues in 2018. Since leaf formation occurs during and immediately following bloom, most of the carbon accumulation required for leaf development would have been either remobilized from stored carbon or acquired before water deficits induced a physiological response (Marchi et al. 2015). Therefore, deficits that were initiated 30 days after full bloom would not have had an appreciable impact on leaf δ^13^C. Both stem and root growth, particularly in non-fruiting trees have been shown to occur during the summer period (Kandiah 1979) therefore carbon accumulated during this period would be more enriched relative to carbon fixed during early spring when water deficits were not applied. Badeck et al. (2005) reported that roots and stems were consistently enriched relative to leaves. However, here, this was only observed when water deficits were applied and were not observed in the control. Cernusak et al. (2009) reviewed underlying reasons why autotrophic tissues are, in general, more depleted than non-photosynthetic tissues and one of the primary hypotheses was differences in timing and conditions during the formation of different organs. When trees were consistently watered throughout the season, no differences in δ^13^C between autotrophic and heterotrophic tissues were observed and that was the case in both 2017 and 2018.

**Figure1.**
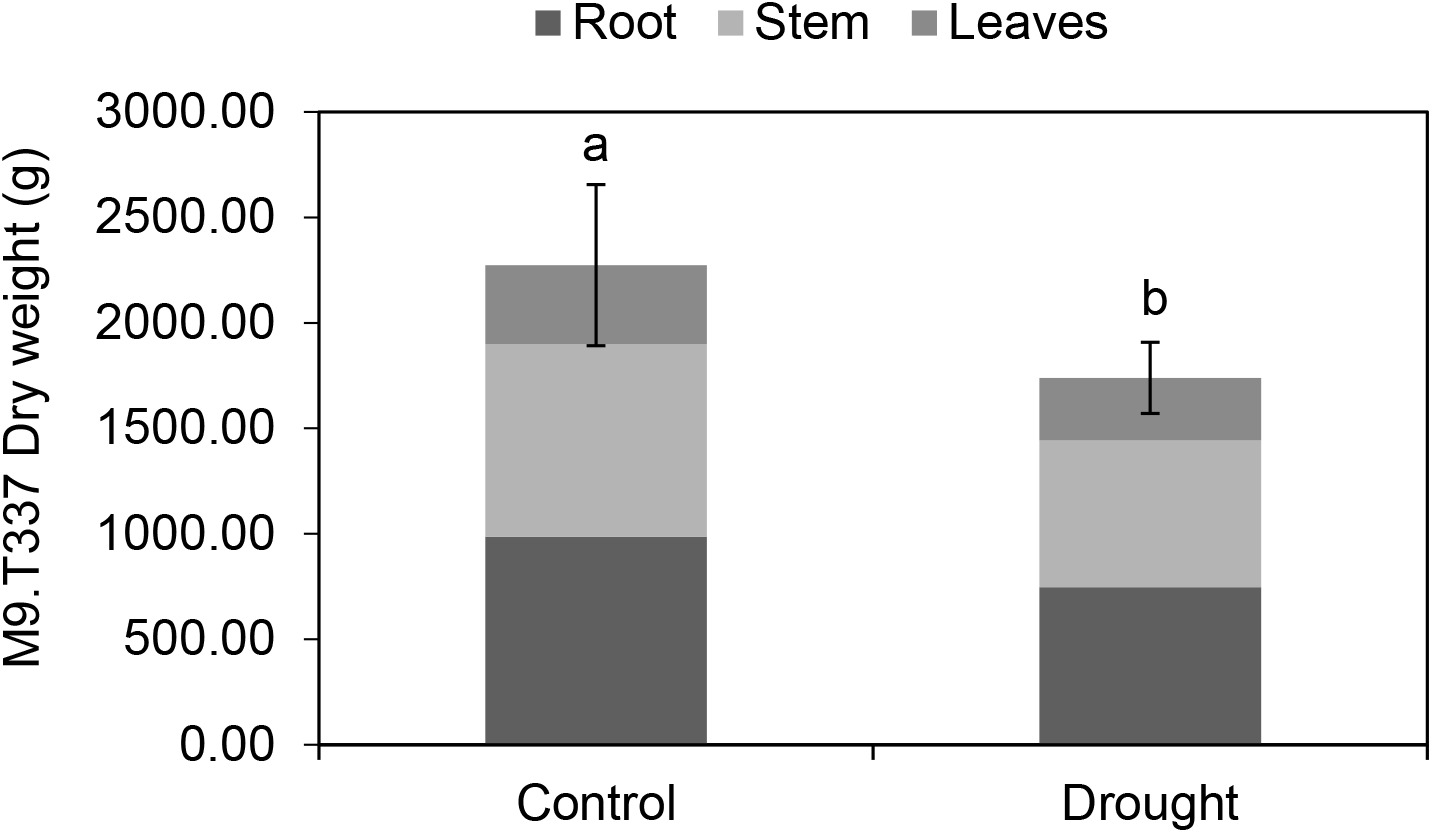
Root, stem and leaf biomass (grams dry weight) of ‘Honeycrisp’ apple trees of 3-year-old grafted on M9-T337 under well-watered control and drought water-stressed treatments for 90 days. Letters denote significant difference between mean total biomass (n=3).

**Figure 2.**
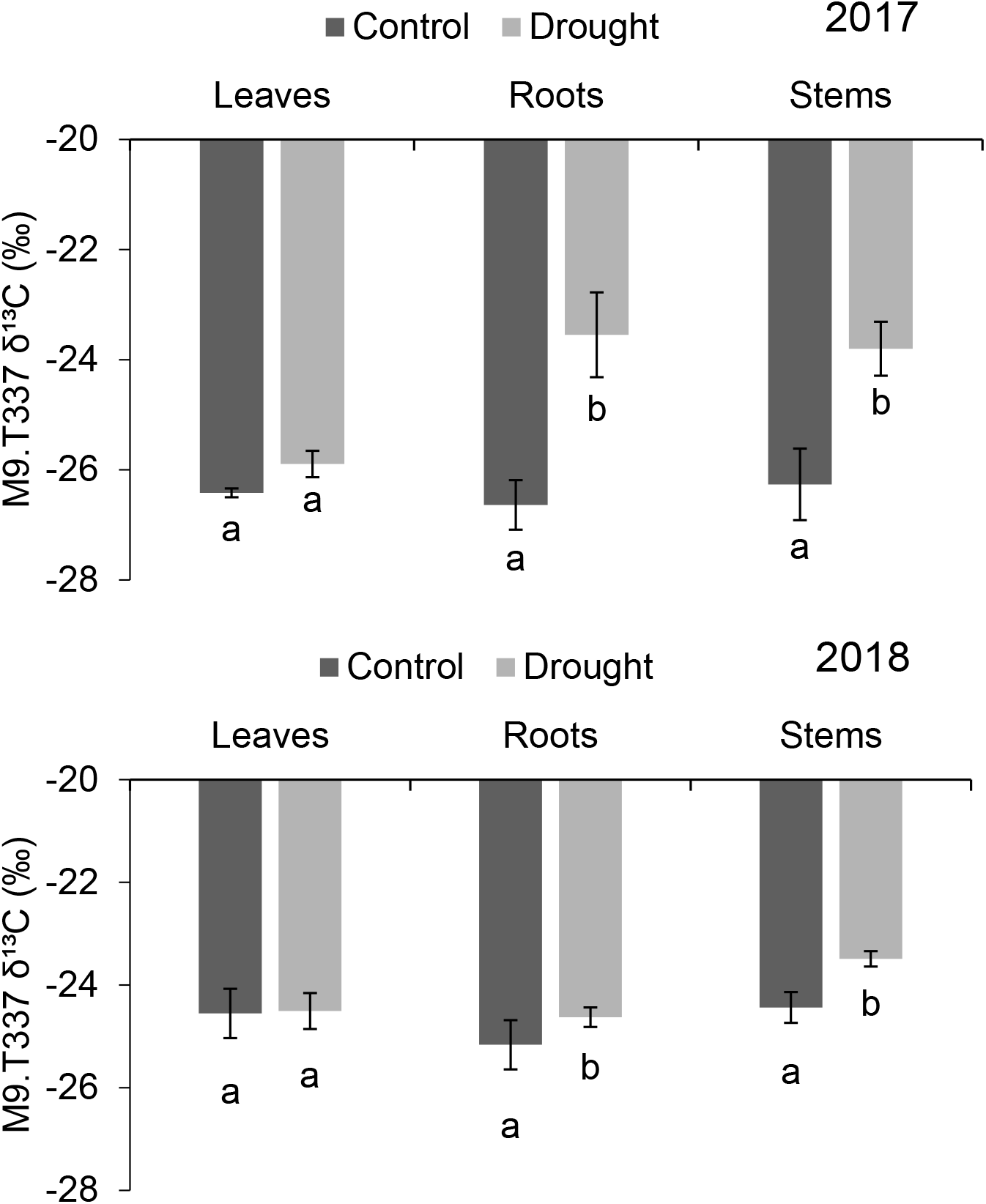
Leaves, roots and stems δ^13^C results for ‘Honeycrisp’ apple trees grafted M9-T337 under well-watered control and drought water-stressed treatments for 90 days in 2017 (top) and 2018 (bottom). Letters denote differences between means determined using a Tukey’s HSD test (n=3).

### Experiment 2

Leaf δ^13^C composition was significantly more depleted in 2017 but fruit δ^13^C was significantly more enriched in 2017 than 2018 (Fig. 3). In both years, fruit were more enriched in δ^13^C relative to leaves and the difference in δ^13^C between fruit and leaves was almost two times greater in 2017 than 2018. Similarly, de Souza et al. (2003) observed strong differences in carbon isotope composition of fruit in grape vines that were water limited to a well-watered control. These results were consistent with the range in differences reported previously for non-photosynthetic tissue compared to leaves (Badeck et al. 2005; Cernusak et al. 2009). Fruit δ^13^C for the well-watered control was significantly more negative (almost 1 ‰) than all other treatments. There were no significant differences in fruit δ^13^C among deficit timing treatments. In 2017, leaf δ^13^C for the early season deficit period was significantly enriched compared to the other treatments. The difference in δ^13^C between leaves and fruit was significantly greater for both late and middle season water limitations than the well-watered control in 2017 but not in 2018.

**Figure 3.**
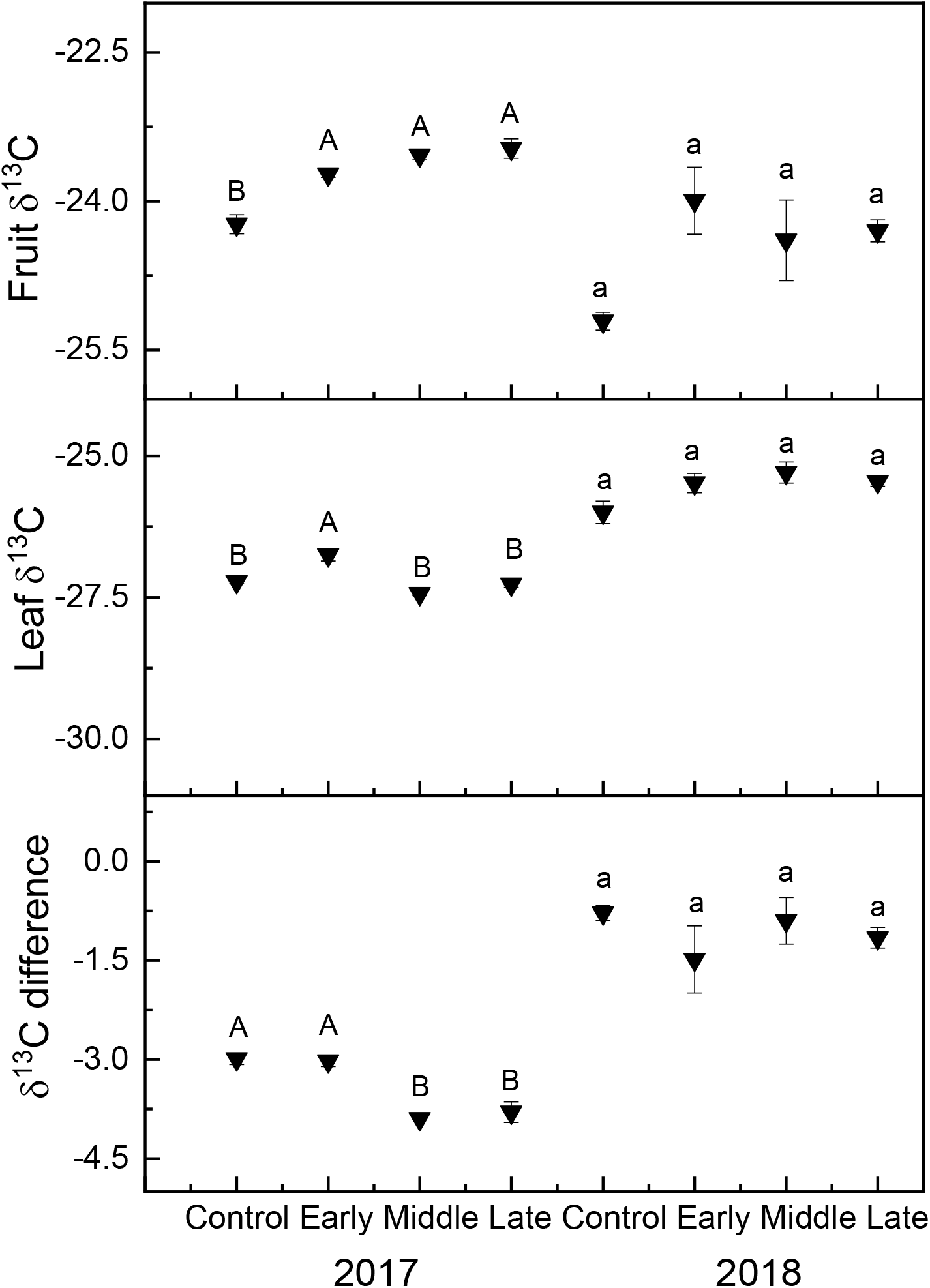
Mean δ^13^C (±SE; N = 3) of ‘Honeycrisp’ apple grafted on M9-T337 under deficit irrigation in fruit (top), leaf (middle), and the difference between leaf and fruit δ^13^Carbon (bottom) in 2017 (left) and 2018 (right). Four treatments were established including a well-watered control, early season deficit (15-45 DAFB), mid-season deficit (45-75 DAFB), and late season deficit (75-105 DAFB; inverted). Uppercase letter denote significance among treatments in 2017 and lowercase letters denote significance among treatments in 2018.

For both experiments, the differences in δ^13^C among treatments were greater in 2017 than in 2018. In 2017, temperatures were warmer, particularly in June and August and smoke present in early August in 2018 reduced transpirational pressure during this time (Table 1). Immediately following bloom, at 0-15 DAFB, mean temperatures were 3.6 °C greater in 2018 than 2017. More importantly, during the mid-season deficit at 46-75 DAFB, mean daily temperatures were 3.2 °C greater in 2017 than in 2018. Temperature is a strong contributor to VPD which, in turn, can strongly affect stomatal conductance (Flore et al. 1984). Stomatal conductance has been closely linked to carbon isotope discrimination in plants (Farquhar et al. 1989; Ehrlinger 1990). This is further supported by more negative δ^13^C values for leaves in 2017 that were reflective of cooler maximum temperatures immediately following bloom. More positive levels in fruit δ^13^C in 2017 were possibly reflective of greater temperatures in June and higher light intensity in August, key periods for carbon loading from leaves to fruit.

**Table 1.** Mean daily temperatures, relative humidity, wind speed, and solar radiation for the Sunrise Research Orchard from May-September in 2017 and 2018.

|  | 2017 |  |  |  |  |
| --- | --- | --- | --- | --- | --- |
|  | Air temp.<br>(°C) | RH<br>(%) | W. Speed<br>(m/s) | Solar rad.<br>(MJ/m²) | Precipitation<br>(mm) |
| May | 16.30 | 51.03 | 1.81 | 22.56 | 21.7 |
| Jun | 20.55 | 40.94 | 2.20 | 24.53 | 6.1 |
| Jul | 26.33 | 29.50 | 2.40 | 27.24 | 0 |
| Aug | 25.02 | 36.06 | 1.82 | 20.59 | 2.2 |
|  | 2018 |  |  |  |  |
| May | 19.88 | 43.64 | 3.16 | 22.49 | 26.8 |
| Jun | 19.93 | 41.02 | 3.01 | 24.58 | 12.0 |
| Jul | 26.01 | 31.54 | 2.89 | 26.37 | 0 |
| Aug | 23.87 | 38.13 | 2.46 | 18.80 | 0.6 |

## CONCLUSION

Leaf carbon isotope composition was a poor indicator of periodic water limitations implemented after bloom in apple. However, δ^13^C for sink tissues that accumulate carbon during deficit periods were much better indicators of water limitations. Interactions with environmental conditions may also contribute to differences between carbon isotope composition of sink and source tissues and physiological response to drought in apple.

## ACKNOWLEDGEMENTS

This work was supported by funding from the Washington Tree Fruit Research Commission (AP-16-101), USDA National Institute of Food and Agriculture - Specialty Crop Research Initiative project “AppleRoot2Fruit: Accelerating the development, evaluation and adoption of new apple rootstocks” (2016-51181-25406), and by the USDA National Institute of Food and Agriculture Hatch project 1014919.

